# TAPAS: Learned integration of AlphaFold3 confidence and geometric features for TCR-pMHC binding prediction

**DOI:** 10.64898/2026.09.08.749548

**Authors:** Ha Young Kim, Hyui Jeong Han, Dongsup Kim

## Abstract

**Motivation:** Recent advances in biomolecular structure prediction, exemplified by AlphaFold3, have opened new opportunities for the prediction of TCR-pMHC binding specificity. Although individual AlphaFold3 confidence metrics provide informative binding signals, their predictive performance varies across datasets, highlighting the need to combine complementary signals rather than rely on any single metric.

**Results:** We present TAPAS, a tabular learning framework that integrates AlphaFold3-derived interface confidence and structural geometry with sequence embeddings. Although no single zero-shot metric was the strongest across all benchmarks, TAPAS was consistently the top-ranked method on VDJdb and two external benchmarks, matching or exceeding the strongest zero-shot AlphaFold3 metric. Feature group ablation showed that the contributions of sequence, confidence, and geometric features varied across evaluation settings, and that combining them resulted in the best overall performance. These results highlight the value of integrating complementary structural and sequence features within a unified framework for robust TCR-pMHC binding prediction.

**Availability and implementation:** Data and source code are available at https://github.com/ha01994/TAPAS.

## 1 Introduction

T cell receptors (TCRs), major components of the adaptive immune system, recognize epitopes presented by major histocompatibility complex (MHC) molecules to trigger immune responses. Predicting interactions between TCRs and peptide-MHC (pMHC) complexes is important for understanding adaptive immunity and for developing treatments for cancer, autoimmune diseases, and infectious diseases. Many prediction tools have been developed, most of which focus on the sequences of TCRs and pMHCs (Feng et al., 2024; Gao et al., 2023; Jensen and Nielsen, 2024; Jiang et al., 2023; Kim et al., 2024; Ye et al., 2025; Zhang et al., 2024). However, predicting TCR-pMHC binding specificity remains a challenging task, particularly for unseen peptides. Recent benchmarking studies further highlight the challenges of generalization in sequence-based TCR-epitope prediction (Drost et al., 2025). Structural information provides a complementary perspective to sequence-based modeling, as TCR-pMHC recognition is ultimately mediated by three-dimensional molecular interactions (McMaster et al., 2024). Driven by advances in computational structure modeling, several studies have attempted to predict TCR-pMHC binding using structure-based approaches (Bradley, 2023; Cao et al., 2026; Deleuran and Nielsen, 2025; Karnaukhov et al., 2024; Siriarchawatana et al., 2026; Slone et al., 2025).

Recent studies have shown that AlphaFold3 (AF3) (Abramson et al., 2024) achieves strong performance in predicting TCR-pMHC structures, highlighting the potential value of structural confidence features derived from predicted complexes (Ascunce-París et al., 2026; Lu et al., 2026). The use of structural confidence metrics is supported by prior work on AlphaFold-based complex prediction, where interface-level metrics such as ipTM, pDockQ, and pDockQ2 have been used to estimate the quality of predicted protein complex interfaces (Bryant et al., 2022; Evans et al., 2021; Zhu et al., 2023). Extending this idea to TCR-pMHC binding, recent studies showed that AlphaFold3-derived confidence metrics are strongly associated with TCR-pMHC binding (Ascunce-París et al., 2026; Messemaker et al., 2025). This view is further supported by IMMREP25, where top-performing methods used zero-shot AlphaFold3 confidence metrics (Richardson et al., 2026). Collectively, these studies indicate that AlphaFold3-derived structural confidence metrics provide informative signals for TCR-pMHC binding prediction.

Despite these advances, recent approaches often rely on individual AlphaFold3 confidence metrics, without fully exploiting the complementary information in sequence, confidence, and geometric features. Moreover, no single metric is reliably best, as performance varies across datasets and evaluation settings. To address these limitations, we propose TAPAS (TCR-Antigen binding Prediction with AlphaFold3 Structural features), a tabular learning framework that combines AlphaFold3-derived interface confidence and structural geometry with ESM-2 sequence embeddings. We evaluated TAPAS on VDJdb and two external benchmarks, ePytope-TCR and IMMREP25, against AlphaFold3-derived zero-shot confidence metrics and sequence-based predictors. Although the best-performing zero-shot metric differed across datasets, TAPAS was consistently the top-ranked method, matching or exceeding the strongest zero-shot AlphaFold3 confidence metric. Rather than relying on any single confidence metric, TAPAS learns to integrate complementary structural and sequence features into a unified predictor, achieving robust performance across datasets.

## 2 Materials and Methods

### 2.1 Dataset

We used VDJdb (Shugay et al., 2018) (downloaded 2025-08-13) as the primary dataset for training and evaluating TAPAS. We restricted the data to human TCRs, HLA class I, paired αβ chains, peptides of length 9-12 and CDR3 sequences of length 8-18. Additionally, we filtered entries to include only those with a VDJdb confidence score greater than zero. This preprocessing resulted in 2,149 TCR-pMHC pairs, spanning 207 unique peptides. All data were partitioned into five folds under random split and strict split settings. In random split, TCR-pMHC pairs were randomly assigned to train, validation, and test sets. In strict split, test peptides were absent from the training set to evaluate generalization to unseen peptides. Under the random split, 80% of the pairs were assigned to the training pool and 20% to the test set. Under the strict split, approximately 80% of the peptides were assigned to the training pool and 20% to the test set. In both settings, 10% of the training pool was reserved for validation.

Negative samples were generated independently within each partition in each fold by random shuffling, which has been reported to reduce negative-sampling bias compared with sampling from a background TCR repertoire (Dens et al., 2023; Moris et al., 2021). For each positive pair, one negative was created by re-pairing its peptide with a TCR originally assigned to a different peptide whose sequence differed from the target peptide by a Levenshtein distance greater than 3, without replacement. This yielded a 1:1 positive-to-negative ratio. Detailed dataset distribution is shown in Supplementary Figure S1 and the cross-validation dataset statistics are shown in Supplementary Table S1.

We used the viral epitope benchmark introduced in the ePytope-TCR study as an independent external evaluation dataset (Drost et al., 2025). The original dataset contains 638 TCRs assigned to 14 viral pMHC targets based on DNA-barcoded pMHC multimer staining. To prevent overlap with the training data, we removed six peptides that overlap with VDJdb and the TCRs assigned to them. This resulted in 445 TCRs and eight pMHC targets. Following the original benchmark, each experimentally assigned TCR-pMHC pair was labeled as positive. Each TCR was then paired with the other seven pMHC targets, and these unassigned pairs were treated as negatives. The resulting dataset contained 3,560 pairs, including 445 positives and 3,115 negatives.

We additionally evaluated TAPAS using the IMMREP25 benchmark dataset (Richardson et al., 2026). IMMREP25 contains 1,000 paired-chain TCRs specific to 20 previously unseen viral peptides, with 10 peptides restricted by HLA-A*02:01 and 10 restricted by HLA-B*40:01. For each peptide, 50 cognate TCRs were labeled as positives, while 450 TCRs assigned to other peptides restricted by the same HLA allele were treated as negatives. The resulting dataset contained 10,000 pairs, including 1,000 positives and 9,000 negatives.

### 2.2 Structure prediction

AlphaFold3 v3.0.3 (Abramson et al., 2024) was run on a single NVIDIA RTX 4500 Ada Generation GPU in order to predict TCR-pMHC structures. Five diffusion samples were generated per pair under default inference settings, and the sample with the highest AlphaFold3 ranking score was retained for TAPAS feature extraction. For negative TCR-pMHC pairs generated in VDJdb, we reused the AlphaFold3 MSAs obtained from positive pairs and re-matched TCRs and pMHCs to construct MSAs for the negative pairs. For the ePytope-TCR and IMMREP25 datasets, MSAs were generated separately for TCRs and MHCs, while peptide MSAs were not generated. To improve computational efficiency, metagenomic and RNA-related databases were excluded during TCR MSA generation. Structure prediction was then performed by matching the pre-computed MSAs for each TCR-pMHC pair.

### 2.3 TAPAS model

We employed TabPFN (Tabular Prior-Data Fitted Network) (Hollmann et al., 2025) as our primary classifier for TCR-pMHC binding prediction (Figure 1). TabPFN is a transformer-based foundation model designed for tabular classification tasks. It is pre-trained on a large number of synthetic tabular datasets and performs in-context learning without requiring additional training or hyperparameter tuning. Given a new dataset, TabPFN predicts class probabilities by conditioning on the training set as context and directly performing inference on the query instances. For our implementation, we used TabPFN v2.5, which supports datasets with up to 50,000 data points and 2,000 features.

**Figure 1.**
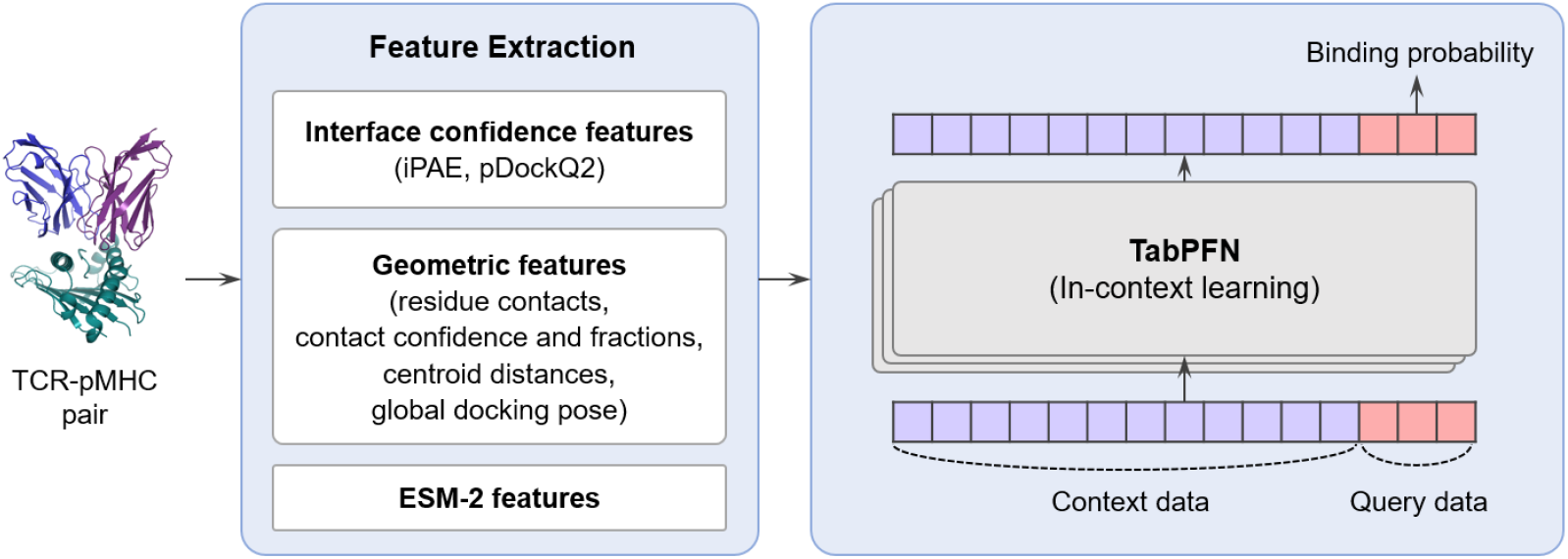
Overview of the TAPAS pipeline for TCR-pMHC binding prediction. Given a TCR-pMHC pair, three types of features are extracted: (i) interface confidence metrics, (ii) geometric features, and (iii) PCA-reduced ESM-2 embeddings of CDR loops and the peptide. These features are concatenated and used as input to TabPFN, which outputs binding probabilities via in-context learning.

Each TCR-pMHC pair was represented by four interface confidence features, 11 geometric features, and 288 ESM-2 sequence features. The confidence features were derived from AlphaFold3-predicted structures and included interface PAE (iPAE) confidence scores and pDockQ2 (Zhu et al., 2023), each computed separately for the TCR and pMHC interfaces. iPAE confidence was calculated for TCR-pMHC interface residue pairs by transforming each PAE value as [1+(PAE/10 Å)^2^]^−1^ and averaging the transformed values. This transformation converts raw PAE into a confidence-like score, with higher values indicating greater confidence in the predicted interface.

The geometric features were calculated from the AlphaFold3-predicted coordinates and included residue contact counts between the peptide and CDR3α, CDR3β, or both CDR3 chains; the fractions of CDR3 and peptide residues involved in these contacts; and confident-contact counts satisfying both pLDDT ≥ 70 and PAE ≤ 10 Å. We additionally calculated centroid distances between the peptide and either the combined CDR3α/β regions or the full TCR, as well as the orientation of the TCR relative to the pMHC complex. Residue contacts were defined using a distance threshold of 5 Å. Details of the geometric features are provided in Supplementary Table S2.

ESM-2 (Lin et al., 2023) 650M model embeddings were used to capture sequence-level information for the peptide and six CDR regions. Residue representations from the final layer were mean-pooled over amino acid tokens, yielding a 1,280-dimensional vector per sequence. Principal component analysis was applied separately to the peptide and CDRs. The peptide, CDR1α, CDR2α, CDR1β, and CDR2β embeddings were each reduced to 32 dimensions, whereas the CDR3α and CDR3β embeddings were each reduced to 64 dimensions, yielding 288 sequence features in total. For VDJdb cross-validation, PCA was fitted on each fold’s training set and applied to its test set. A validation partition was available but not used by TAPAS, since all settings were fixed and no tuning or model selection was performed. The validation partitions were used only for checkpoint selection in the trainable sequence-based baselines.

For the external ePytope-TCR and IMMREP25 benchmarks, we constructed a 10-model ensemble comprising one model per cross-validation fold, corresponding to five random split and five strict split folds. Each model was trained on all 4,298 TCR-pMHC pairs of its fold, pooling the training, validation, and test partitions. Because negatives were sampled independently per fold, the ten models differed in their negative pairings, providing ensemble diversity. For each ensemble member, PCA was fitted on the full 4,298-pair VDJdb set and applied to the external data. The final binding score was obtained by averaging the predicted probabilities across the 10 models. Across the external benchmarks, this ensemble exceeded the average performance of the individual fold models while yielding more stable predictions (Supplementary Figure S2).

### 2.4 Baseline methods

We used four AlphaFold3 confidence metrics as zero-shot predictors, each computed from the structure with the highest AF3 ranking score: iPAE confidence and pDockQ2, each averaged over the TCR chains and over the pMHC chains and then averaged across the two sides; TCR-pMHC ipTM, the mean of the four chain-pair ipTM values (MHC-TCRα, MHC-TCRβ, peptide-TCRα, peptide-TCRβ); and pLDDT, the mean per-atom pLDDT over the entire complex. We also compared TAPAS to sequence-based models, TSpred (Kim et al., 2024) and NetTCR-2.2 (Jensen and Nielsen, 2024). Both models were retrained on the same VDJdb splits as TAPAS to ensure a fair comparison. For the external benchmarks, both were applied as the same 10-model ensemble used for TAPAS.

### 2.5 Evaluation and statistical analysis

Following IMMREP25, we adopted macro-AUC@0.1 as the primary evaluation metric, defined as the partial area under the ROC curve restricted to the region FPR ≤ 0.1 and macro-averaged across peptides. This metric focuses on performance when only a small number of false positives are allowed, which is important in TCR specificity prediction where most candidates are non-binders and the goal is to identify a small number of true binders. Partial AUC values were standardized using scikit-learn’s roc_auc_score with max_fpr=0.1. We additionally report the full macro-AUC as a secondary metric in Supplementary Table S3. Because VDJdb contains many singleton and low-frequency peptides, many peptide-level evaluation groups had only a few negative examples. In these small groups, the standardized partial AUC used by scikit-learn can have an expected random value above 0.5. Under our evaluation procedure, the expected macro-AUC@0.1 for random rankings without ties was approximately 0.6864 for random split and 0.6780 for strict split. We therefore interpret the results together with full macro-AUC (Supplementary Table S3).

For each dataset, TAPAS was compared against the best-performing confidence metric on that dataset. All statistical uncertainty was assessed at the peptide level, treating each peptide as the resampling unit. To obtain confidence intervals, we generated 10,000 paired bootstrap replicates: in each replicate, peptides were resampled with replacement while keeping each method’s predictions paired within a peptide. The 95% confidence interval for the difference in macro-AUC@0.1 between two methods was taken as the 2.5th and 97.5th percentiles of these replicates. To test significance, we used a two-sided paired sign-flip permutation test: under the null hypothesis of no systematic difference between two methods, the sign of each peptide-specific performance difference was randomly flipped, and the resulting null distribution was used to compute p-values. For VDJdb, peptide-specific AUC@0.1 values were first averaged within each test fold and then across the five folds. In the random split, results for the same peptide across different folds were grouped together in the bootstrap and sign-flip tests.

For the VDJdb strict split, we examined train-test peptide sequence similarity within each fold. For each test peptide, we identified the most similar peptide in the training set using normalized Levenshtein identity, defined as 1 - d_Lev(x,y)/max(|x|,|y|), where d_Lev is the minimum number of amino acid substitutions, insertions, or deletions required to transform one sequence into the other. As a sensitivity analysis, we excluded test peptides sharing at least 80% identity with any training peptide and recomputed macro-AUC@0.1 for TAPAS and the strongest zero-shot metric.

## 3 Results

### 3.1 Results on VDJdb random split and strict split

We evaluated TAPAS on VDJdb random split and strict split, comparing against several zero-shot metrics derived from AlphaFold3 predictions as well as sequence-based methods (Table 1). TAPAS achieved a macro-AUC@0.1 of 0.8985 under the random split, outperforming all evaluated AlphaFold3-derived zero-shot metrics and sequence-based models. Compared with TCR-pMHC ipTM, the best-performing zero-shot metric on this dataset, TAPAS showed a paired improvement of +0.0592 (95% peptide-level paired-bootstrap CI, 0.0321 to 0.0863; two-sided paired sign-flip p<0.001). Under the strict split, TAPAS achieved a macro-AUC@0.1 of 0.8425. Its performance was numerically higher than that of iPAE confidence, the strongest zero-shot metric on this dataset, but the paired difference was small (Δ=+0.0054; 95% peptide-level paired-bootstrap CI, -0.0190 to 0.0304; two-sided paired sign-flip p=0.671). Similarly, TAPAS achieved the numerically highest performance in terms of full macro-AUC under both splits, but the difference was small for the strict split (Supplementary Table S3). These results indicate that TAPAS provides a clear advantage when peptides are represented during training, whereas its additional improvement over the strongest AlphaFold3 confidence metric was small for unseen peptides. Notably, the sequence-based models dropped sharply under the strict split, far below all confidence metrics.

**Table 1.** Performance of TAPAS and other methods in terms of macro-AUC@0.1 on the VDJdb random split and strict split cross-validation datasets. Values are the mean macro-AUC@0.1 across the five folds.

| Method | Random split<br>macro-AUC@0.1 | Strict split<br>macro-AUC@0.1 |
| --- | --- | --- |
| TAPAS | **0.8985** | **0.8425** |
| Zero-shot iPAE confidence | 0.8390 | 0.8371 |
| Zero-shot pDockQ2 | 0.8309 | 0.8333 |
| Zero-shot TCR-pMHC ipTM | 0.8393 | 0.8281 |
| Zero-shot pLDDT | 0.8178 | 0.8006 |
| TSpred | 0.8311 | 0.7170 |
| NetTCR-2.2 | 0.8174 | 0.7171 |

To assess whether the strict split performance depended on peptides similar to the training set, we performed a sensitivity analysis excluding test peptides with at least 80% identity to any training peptide. After this exclusion, TAPAS retained a macro-AUC@0.1 of 0.8338, compared with 0.8254 for iPAE confidence, showing that the difference between the two methods was not reduced by this exclusion (Supplementary Table S4).

### 3.2 Results on unseen epitopes in the ePytope-TCR benchmark

We further assessed the performance of TAPAS ensemble on the ePytope-TCR viral epitope benchmark (Table 2). The TAPAS ensemble achieved a macro-AUC@0.1 of 0.5618, compared with 0.5585 for zero-shot TCR-pMHC ipTM, the strongest zero-shot metric on this benchmark. TAPAS therefore achieved numerically higher performance, although the difference was small (Δ=+0.0033; 95% peptide-level paired-bootstrap CI, -0.0136 to 0.0205; two-sided paired sign-flip p=0.734). As under the strict split, the sequence-based models performed close to random, whereas all AlphaFold3-based approaches retained a modest but consistent predictive signal. Under the full macro-AUC (Supplementary Table S3), the zero-shot confidence metrics were marginally higher than TAPAS on ePytope-TCR, in contrast to the other benchmarks where TAPAS remained highest.

**Table 2.** Performance of TAPAS and other methods in terms of macro-AUC@0.1 on the ePytope-TCR viral epitope benchmark.

| Method | macro-AUC@0.1 |
| --- | --- |
| TAPAS ensemble | **0.5618** |
| Zero-shot iPAE confidence | 0.5540 |
| Zero-shot pDockQ2 | 0.5547 |
| Zero-shot TCR-pMHC ipTM | 0.5585 |
| Zero-shot pLDDT | 0.5545 |
| TSpred | 0.5087 |
| NetTCR-2.2 | 0.5041 |

### 3.3 Results on unseen epitopes in the IMMREP25 benchmark

We additionally evaluated TAPAS ensemble on the unseen peptides of the IMMREP25 benchmark dataset (Table 3). On this benchmark, the winning submission used a zero-shot AlphaFold-based pLDDT score. Because the official challenge score was calculated on an unreleased private-leaderboard subset, we did not directly compare it with our results obtained on the complete released dataset. As shown in Table 3, the TAPAS ensemble achieved a macro-AUC@0.1 of 0.5847, outperforming zero-shot pDockQ2, the best-performing zero-shot metric on this benchmark (Δ=+0.0107; 95% peptide-level paired-bootstrap CI, 0.0013 to 0.0235; two-sided paired sign-flip p=0.047). As in the other benchmarks, the sequence-based models performed close to random. We further examined performance at the peptide level on both external benchmarks (Supplementary Figure S3). TAPAS outperformed the strongest zero-shot metric for four of eight ePytope-TCR peptides, and for 11 of 20 IMMREP25 peptides.

**Table 3.** Performance of TAPAS and other methods in terms of macro-AUC@0.1 on the IMMREP25 dataset.

| Method | macro-AUC@0.1 |
| --- | --- |
| TAPAS ensemble | **0.5847** |
| Zero-shot iPAE confidence | 0.5722 |
| Zero-shot pDockQ2 | 0.5740 |
| Zero-shot TCR-pMHC ipTM | 0.5732 |
| Zero-shot pLDDT | 0.5600 |
| TSpred | 0.5078 |
| NetTCR-2.2 | 0.5002 |

As a sensitivity analysis, we applied to all methods the score normalization and cluster smoothing procedure used by the top-ranked submission in IMMREP25 (Richardson et al., 2026). This procedure subtracts peptide- and TCR-specific mean scores and then applies TCRdist-based single-linkage cluster smoothing (Supplementary Note S1). TAPAS remained the highest-performing method at each processing stage, with macro-AUC@0.1 increasing from 0.5847 for the raw scores to 0.6020 after normalization and smoothing (Supplementary Table S5).

### 3.4 Ablation studies and feature interpretation

To assess the contribution of the different feature groups, we conducted an ablation study on the VDJdb dataset using various feature combinations (Table 4). The three feature groups showed markedly different behavior across the two splits. Under the random split, ESM-2 embeddings alone achieved the highest single-group performance, substantially exceeding the confidence-only and geometry-only models, indicating that sequence information is the dominant signal when peptides are seen during training. However, under the strict split, the performance of ESM-2 alone dropped sharply, whereas the confidence-only and geometry-only models were more robust. The effects of combining feature groups varied across combinations and splits. Nevertheless, the full TAPAS model achieved the highest performance under both splits. Its improvement over the best-performing single feature group was larger under the strict split, suggesting that feature integration is particularly beneficial for unseen peptides. The full model also performed best on the external benchmarks (Supplementary Table S6).

**Table 4.** Feature ablation on VDJdb. The three feature groups are ESM-2 sequence embeddings, AlphaFold3 confidence features, and geometric features. The full TAPAS model integrates all three groups. Values are the mean macro-AUC@0.1 across the five folds.

| Method | Random split<br>macro-AUC@0.1 | Strict split<br>macro-AUC@0.1 |
| --- | --- | --- |
| ESM-2 only | 0.8634 | 0.7246 |
| Confidence only | 0.8314 | 0.7755 |
| Geometry only | 0.7783 | 0.7736 |
| ESM-2 + confidence | 0.8928 | 0.8327 |
| ESM-2 + geometry | 0.8946 | 0.8161 |
| Confidence + geometry | 0.8094 | 0.7870 |
| Full TAPAS | **0.8985** | **0.8425** |

To justify our choice of classifier, we compared TabPFN against several standard classifiers trained on the same TAPAS feature set: random forest, XGBoost, a multilayer perceptron, and logistic regression (Table 5). TabPFN achieved the best macro-AUC@0.1 under both splits. TabPFN performed comparably to random forest under the random split but showed a clearer advantage under the strict split. It also achieved the highest numerical performance on both external benchmarks (Supplementary Table S7). Additionally, using IMMREP25 as a representative benchmark, we profiled the computational cost of the pipeline. AF3 structure generation was the dominant cost, requiring approximately 780 s per pair compared with 12 s per pair for downstream feature extraction, while TAPAS fitting and inference across all 10 ensemble models required 151 s in total (Supplementary Table S8).

**Table 5.** Comparison of classifiers using the TAPAS feature set on VDJdb. Values are the mean macro-AUC@0.1 across the five folds.

| Model | Random split<br>macro-AUC@0.1 | Strict split<br>macro-AUC@0.1 |
| --- | --- | --- |
| TabPFN | **0.8985** | **0.8425** |
| Random forest | 0.8955 | 0.8197 |
| XGBoost | 0.8701 | 0.8125 |
| MLP | 0.8496 | 0.7954 |
| Logistic regression | 0.7820 | 0.7661 |

We further applied SHAP (SHapley Additive exPlanations) analysis (Lundberg and Lee, 2017) to interpret the contribution of structural features in TAPAS (Supplementary Note S2). As permutation SHAP over the full feature set was computationally prohibitive, we performed this analysis on a structure-only submodel trained on the four confidence and 11 geometric features, excluding the 288-dimensional ESM-2 features. The SHAP results below therefore describe feature importance within this structural submodel rather than the full TAPAS model. SHAP analyses were conducted in four evaluation settings: the VDJdb random split and strict split fold-0 models on their respective held-out test sets, and a model trained on the complete VDJdb random split fold-0 data and applied to IMMREP25 and ePytope-TCR.

Mean absolute SHAP values were normalized within each dataset to sum to one, yielding each feature’s relative contribution (Figure 2). Across all four settings, pDockQ2 interface scores accounted for the largest contribution. The pMHC-interface pDockQ2 score ranked first in every setting, followed by the TCR-interface pDockQ2 score, while iPAE confidence scores contributed next and geometric features showed smaller and more variable relative contributions. Signed SHAP values (Supplementary Figure S4) indicated that higher pDockQ2 values consistently shifted predictions toward binding. Within the structural submodel, prediction relied primarily on AlphaFold3 interface confidence signals, particularly pDockQ2, supplemented by geometric information.

**Figure 2.**
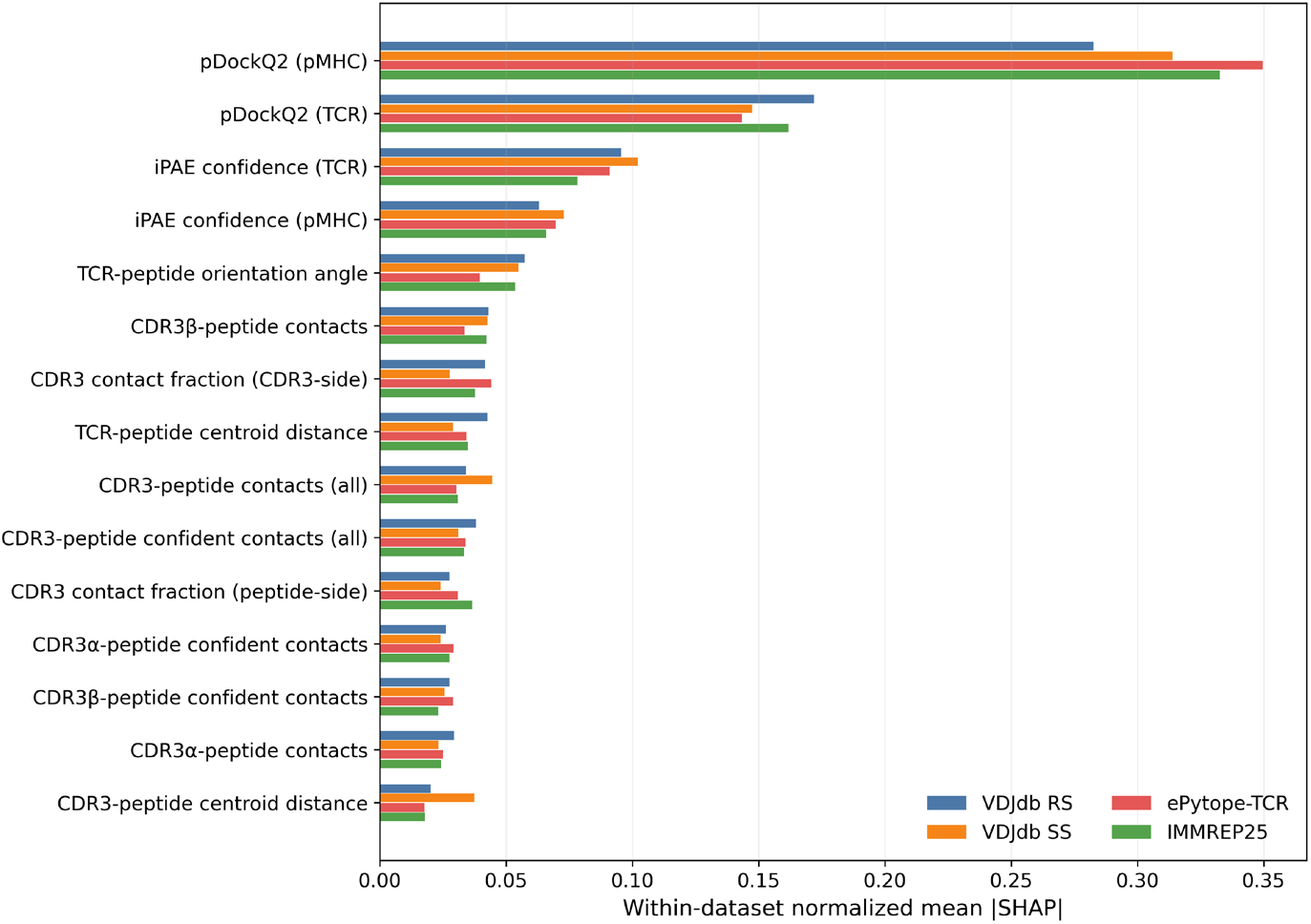
Relative SHAP importance of structural features across evaluation settings in the structure-only submodel. For each of the 15 structural features, bars show the within-dataset normalized mean absolute SHAP value. Colors denote the four evaluation settings: VDJdb RS (random split), VDJdb SS (strict split), ePytope-TCR, and IMMREP25.

## 4 Discussion

In this study, we proposed TAPAS, a tabular learning framework that integrates AlphaFold3-derived interface confidence, geometric features and sequence embeddings for TCR-pMHC binding prediction. Across VDJdb and two external benchmarks, TAPAS performed comparably to or better than the strongest zero-shot confidence metrics. Feature ablation further showed that the contribution of each feature group depended on the evaluation setting, with the integrated model performing best overall. These results indicate that integrating complementary structural and sequence features within a unified framework provides consistent performance across diverse evaluation settings.

A central observation across our benchmarks is that no single zero-shot confidence metric was consistently the strongest: TCR-pMHC ipTM was best under the VDJdb random split and on ePytope-TCR, iPAE under the strict split, and pDockQ2 on IMMREP25. In practice, this means that selecting the optimal zero-shot metric for a new dataset is not straightforward, as the best-performing metric cannot be known in advance. By learning to integrate multiple confidence, geometric, and sequence features, TAPAS matched or exceeded the strongest zero-shot metric in every setting without requiring such a choice.

In our comparison with conventional classifiers trained on the same feature set, TabPFN performed better on unseen peptides, suggesting improved generalization of TabPFN to out-of-distribution peptides. Moreover, because it requires no gradient-based task-specific retraining, TabPFN can be readily applied to new datasets once the corresponding structures are available. Its tabular framework also allows new features to be incorporated in a straightforward manner, enabling continuous improvement as more complementary data sources become available.

Despite these improvements, several limitations remain. While AlphaFold3 confidence metrics provide informative signals, they depend on the quality of predicted structures. As a result, the model may not fully capture all types of TCR-pMHC interactions, especially those that are underrepresented in available structural data. In addition, TAPAS currently relies on a single predicted structure selected by the highest AF3 ranking score, even though prediction accuracy can depend on sampling and model selection strategies (Elofsson, 2026). Future work may address these limitations by improving structural modeling accuracy and incorporating multiple predicted structures per TCR-pMHC pair to enhance robustness.

Recent work (Wang et al., 2026) has demonstrated that deep peptide recognition profiling (PRP), combined with protein language models, can achieve highly accurate prediction of TCR specificity. However, such approaches rely on large-scale experimental profiling, such as yeast display, and have been evaluated in relatively limited experimental settings within a single HLA context. In contrast, once trained, TAPAS uses AlphaFold3-derived structural features to predict new TCR-pMHC pairs without requiring target-specific experimental profiling, while maintaining robust performance across diverse datasets and evaluation settings. These complementary strengths suggest that integrating PRP-derived functional signals with structure-based features may further improve predictive performance in future studies.

## Data Availability

Data and source code are available at https://github.com/ha01994/TAPAS.

