## Supplementary file for "TAPAS: Learned integration of AlphaFold3 confidence and geometric features for TCR-pMHC binding prediction"

**Supplementary Note S1. Score normalization and TCRdist-based smoothing for IMMREP25**

We evaluated a label-free postprocessing procedure based on the score normalization and TCR clustering strategy described in IMMREP25 (Richardson et al., 2026). For each MHC context, predictions were arranged as a score matrix over TCRs (rows) and candidate peptides (columns). In the normalization step, we subtracted the corresponding row mean and column mean from each matrix entry, removing systematic score differences associated with individual TCRs and peptides. The TCRs within each allele were subsequently grouped by single-linkage clustering using TCRdist with a distance threshold of 120. Scores within each cluster were averaged, with each cluster weighted by the square root of its size. Averaging across sequence-similar TCRs reduces variation among individual predictions, whereas square-root scaling partially offsets the reduction in score magnitude caused by averaging. The same normalization and smoothing were applied to all methods without method-specific tuning. Performance at each stage is reported in Supplementary Table S5.

**Supplementary Note S2. Details on SHAP analysis**

Model interpretation was performed on a structural-feature submodel comprising 15 features: 4 interface confidence features and 11 geometric features. The 288-dimensional ESM-2 embedding block was excluded, as applying permutation SHAP to the full feature set was computationally prohibitive. SHAP values were computed using the permutation explainer across four evaluation settings. For VDJdb, values were calculated for the random split (RS) and strict split (SS) fold-0 models using their respective held-out test sets. For the external benchmarks, a model trained on the complete VDJdb RS fold-0 data was applied to ePytope-TCR and IMMREP25.

From each evaluation set, we sampled 100 positive and 100 negative pairs without replacement, and used 100 randomly selected training pairs as the background distribution. Each SHAP value was estimated using 31 model evaluations per sample (2 × number of features + 1), yielding approximately 6,200 TabPFN inference calls per setting. For descriptive comparison of relative feature importance across settings, mean absolute SHAP values were normalized within each dataset to sum to one. These normalized shares describe the relative importance of features within each dataset, and were not used to compare absolute SHAP magnitudes across datasets. Raw signed SHAP values were additionally visualized as beeswarm plots (Supplementary Figure S4).

**Supplementary Figure S1. Distribution of peptide and MHC counts in the VDJdb dataset.** (Top) Peptide frequency distribution shown on a logarithmic scale. 72 peptides with only a single observation were excluded for visualization clarity. (Bottom) MHC allele frequency distribution (log scale).

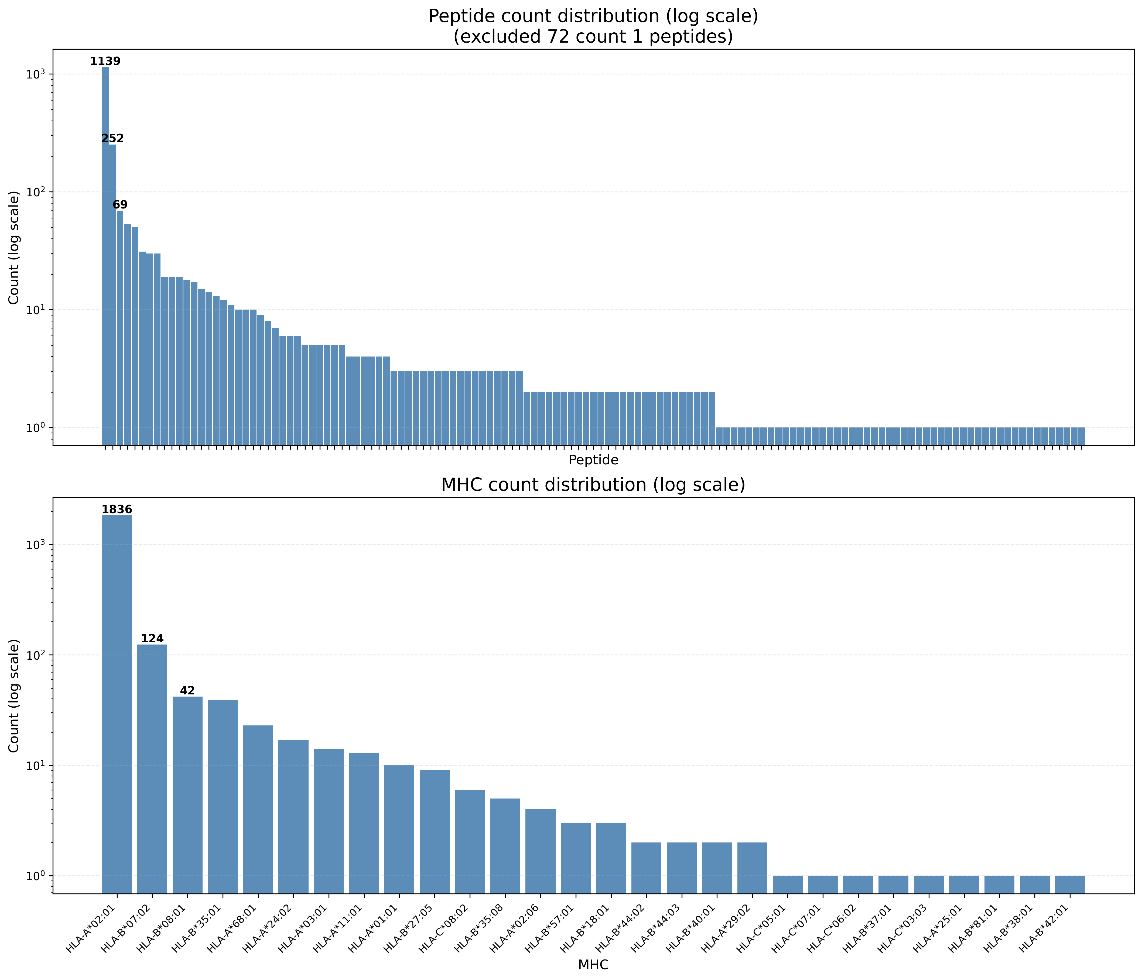

**Supplementary Figure S2. Stability of the TAPAS ensemble across external evaluation datasets.** (A, B) macro-AUC@0.1 values obtained from the 10 individual VDJdb-trained models and their ensemble on (A) ePytope-TCR and (B) IMMREP25. Each point represents an individual model, the red diamond denotes the 10-model ensemble, and the dashed line indicates the mean of the individual models. (C, D) Performance as a function of ensemble size on (C) ePytope-TCR and (D) IMMREP25. Boxplots summarize all model subsets of each ensemble size, the blue line indicates the mean, and the red diamond denotes the full 10-model ensemble.

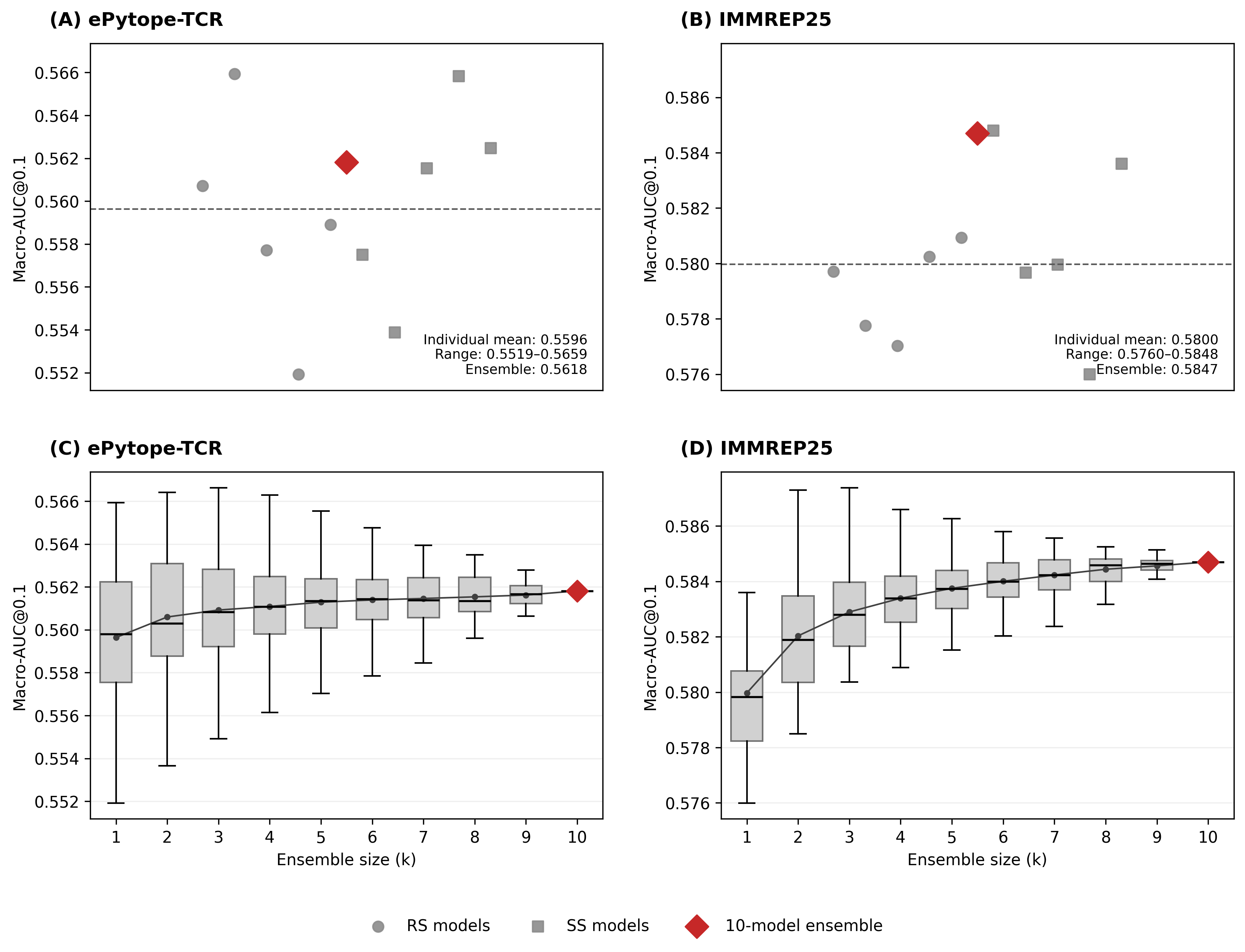

**Supplementary Figure S3. Peptide-level comparison of TAPAS and the strongest zero-shot comparator on the external benchmarks.** Each point represents one peptide, with its per-peptide AUC@0.1 for the TAPAS ensemble (y-axis) plotted against the corresponding zero-shot comparator (x-axis) on (a) ePytope-TCR and (b) IMMREP25. The dashed line denotes equal performance.

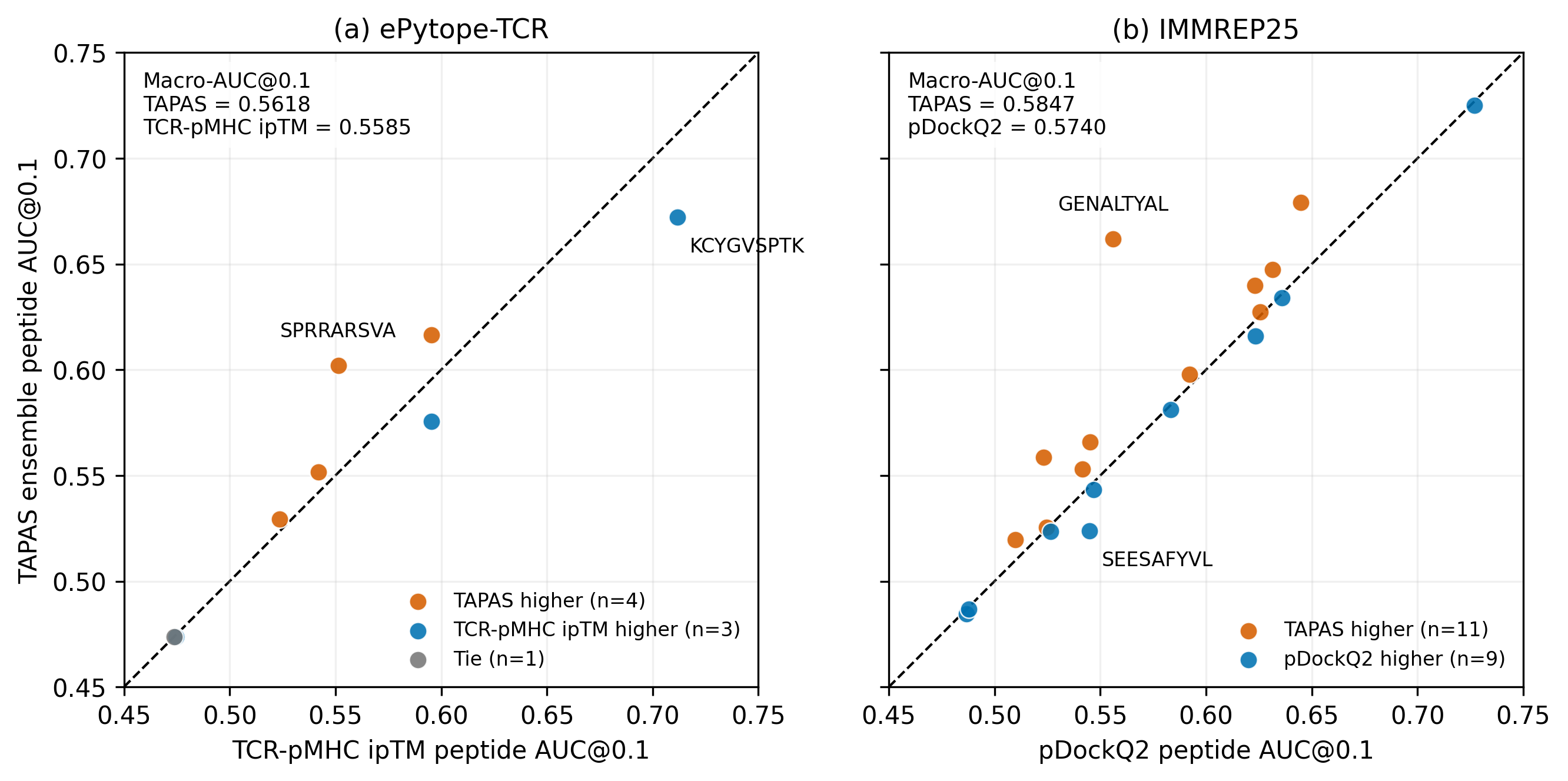

**Supplementary Figure S4. Signed SHAP values for the structure-only submodel across the four evaluation settings.** Beeswarm plots show the distribution of signed SHAP values for the 15 structural features (4 confidence and 11 geometric). Each point represents one sampled test pair and its horizontal position is the feature’s SHAP value for that prediction. Positive values push the prediction toward binding and negative values toward non-binding. Point color encodes the feature value, from low (blue) to high (red). (A) VDJdb random split (fold 0), (B) VDJdb strict split (fold 0), (C) ePytope-TCR, and (D) IMMREP25. Only the top 12 features are shown in each panel.

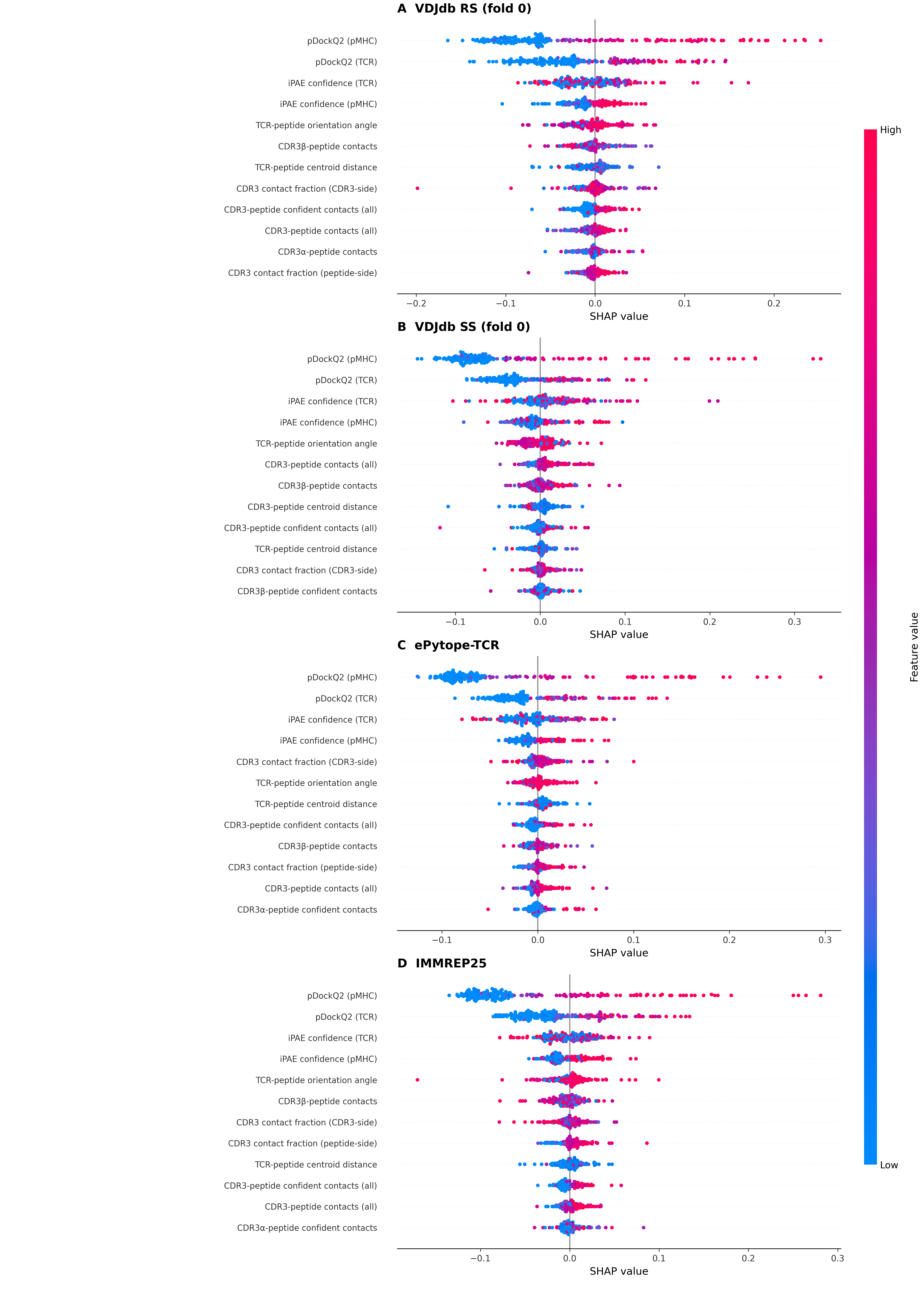

**Supplementary Table S1. Cross-validation fold statistics for VDJdb dataset.** Five-fold cross-validation statistics are shown for random split (RS) and strict split (SS).

| VDJdb random split | | | | | | |
| --- | --- | --- | --- | --- | --- | --- |
| Fold | Train | Val | Test | # Train  peptides | # Val  peptides | # Test  peptides |
| 0 | 3094 | 344 | 860 | 171 | 42 | 68 |
| 1 | 3094 | 344 | 860 | 163 | 40 | 82 |
| 2 | 3094 | 344 | 860 | 167 | 41 | 75 |
| 3 | 3094 | 344 | 860 | 172 | 41 | 67 |
| 4 | 3096 | 344 | 858 | 168 | 35 | 78 |
| VDJdb strict split | | | | | | |
| Fold | Train | Val | Test | # Train  peptides | # Val  peptides | # Test  peptides |
| 0 | 4004 | 88 | 206 | 148 | 17 | 42 |
| 1 | 3164 | 244 | 890 | 148 | 17 | 42 |
| 2 | 3716 | 124 | 458 | 149 | 17 | 41 |
| 3 | 3884 | 130 | 284 | 149 | 17 | 41 |
| 4 | 1764 | 74 | 2460 | 149 | 17 | 41 |

**Supplementary Table S2. Detailed information on the geometric features.** The definitions of all 11 geometric features are listed.

| Feature | Definition |
| --- | --- |
| CDR3α-peptide contacts | Number of CDR3α-peptide residue pairs with any interatomic distance ≤5 Å |
| CDR3β-peptide contacts | Number of CDR3β-peptide residue pairs with any interatomic distance ≤5 Å |
| CDR3-peptide contacts (all) | Number of combined CDR3α/β-peptide residue pairs with any interatomic distance ≤5 Å |
| CDR3 contact fraction (CDR3-side) | Fraction of combined CDR3α/β residues contacting at least one peptide residue within 5 Å |
| CDR3 contact fraction (peptide-side) | Fraction of peptide residues contacting at least one combined CDR3α/β residue within 5 Å |
| CDR3α-peptide confident contacts | Number of CDR3α-peptide residue pairs within 5 Å with mean residue-pair pLDDT ≥70 and symmetrized PAE ≤10 Å |
| CDR3β-peptide confident contacts | Number of CDR3β-peptide residue pairs within 5 Å with mean residue-pair pLDDT ≥70 and symmetrized PAE ≤10 Å |
| CDR3-peptide confident contacts (all) | Number of combined CDR3α/β-peptide residue pairs within 5 Å with mean residue-pair pLDDT ≥70 and symmetrized PAE ≤10 Å |
| CDR3-peptide centroid distance | Euclidean distance between the all-atom centroids of the combined CDR3α/β loops and peptide |
| TCR-peptide centroid distance | Euclidean distance between the all-atom centroids of the full TCR and peptide |
| TCR-peptide orientation angle | Angle in degrees subtended at the peptide centroid by the TCR and MHC centroids |

**Supplementary Table S3. Performance of TAPAS and other methods in terms of macro-AUC across the three benchmarks.** Performance is reported as the macro-averaged AUC computed over the full ROC curve.

a. VDJdb

| Method | Random split  macro-AUC | Strict split  macro-AUC |
| --- | --- | --- |
| TAPAS | **0.8668** | **0.7748** |
| Zero-shot iPAE confidence | 0.7802 | 0.7736 |
| Zero-shot pDockQ2 | 0.7616 | 0.7669 |
| Zero-shot TCR-pMHC ipTM | 0.7756 | 0.7604 |
| Zero-shot pLDDT | 0.7333 | 0.7133 |
| TSpred | 0.7663 | 0.5723 |
| NetTCR-2.2 | 0.7503 | 0.5681 |

b. ePytope-TCR

| Method | macro-AUC |
| --- | --- |
| TAPAS ensemble | 0.6019 |
| Zero-shot iPAE confidence | 0.6214 |
| Zero-shot pDockQ2 | 0.6160 |
| Zero-shot TCR-pMHC ipTM | 0.6167 |
| Zero-shot pLDDT | **0.6238** |
| TSpred | 0.5414 |
| NetTCR-2.2 | 0.5354 |

c. IMMREP25

| Method | macro-AUC |
| --- | --- |
| TAPAS ensemble | **0.6390** |
| Zero-shot iPAE confidence | 0.6349 |
| Zero-shot pDockQ2 | 0.6334 |
| Zero-shot TCR-pMHC ipTM | 0.6315 |
| Zero-shot pLDDT | 0.6234 |
| TSpred | 0.5344 |
| NetTCR-2.2 | 0.5088 |

**Supplementary Table S4. Remote-subset sensitivity analysis on the VDJdb strict split.** As a sensitivity analysis, macro-AUC@0.1 was recomputed after excluding test peptides sharing at least 80% identity with any training peptide. Values are the mean macro-AUC@0.1 across the five folds; iPAE confidence is shown as the strongest zero-shot metric on this split. Removing near-duplicate peptides produced only a modest decrease for both methods and did not reduce the numerical difference between TAPAS and iPAE confidence.

| Evaluation subset | Peptides | Pairs | TAPAS | iPAE confidence | TAPAS - iPAE |
| --- | --- | --- | --- | --- | --- |
| Full strict split | 207 | 4,298 | 0.8425 | 0.8371 | +0.0054 |
| Excluding identity ≥ 80% | 167 | 1,758 | 0.8338 | 0.8254 | +0.0084 |

**Supplementary Table S5. Effect of postprocessing on the IMMREP25 benchmark.** Macro-AUC@0.1 for each method at three stages: raw scores (Raw), after normalization (Normalization), and after normalization followed by TCRdist single-linkage cluster smoothing (Normalization + smoothing).

| Method | macro-AUC@0.1  (Raw) | macro-AUC@0.1  (Normalization) | macro-AUC@0.1  (Normalization + smoothing) |
| --- | --- | --- | --- |
| TAPAS ensemble | **0.5847** | **0.5939** | **0.6020** |
| Zero-shot iPAE confidence | 0.5722 | 0.5700 | 0.5746 |
| Zero-shot pDockQ2 | 0.5740 | 0.5911 | 0.5948 |
| Zero-shot TCR-pMHC ipTM | 0.5732 | 0.5704 | 0.5738 |
| Zero-shot pLDDT | 0.5600 | 0.5734 | 0.5766 |
| TSpred | 0.5078 | 0.5141 | 0.5287 |
| NetTCR-2.2 | 0.5002 | 0.5077 | 0.5096 |

**Supplementary Table S6. Feature ablation on the external benchmarks (ePytope-TCR, IMMREP25).** The three feature groups are ESM-2 sequence embeddings, AlphaFold3 confidence features, and geometric features. The full TAPAS model integrates all three groups. Each value is the macro-AUC@0.1 of the corresponding 10-model ensemble.

a. ePytope-TCR

| Feature set | macro-AUC@0.1 |
| --- | --- |
| ESM-2 only | 0.5013 |
| Confidence only | 0.5434 |
| Geometry only | 0.5372 |
| ESM-2 + confidence | 0.5608 |
| ESM-2 + geometry | 0.5431 |
| Confidence + geometry | 0.5566 |
| Full TAPAS | **0.5618** |

b. IMMREP25

| Feature set | macro-AUC@0.1 |
| --- | --- |
| ESM-2 only | 0.5009 |
| Confidence only | 0.5757 |
| Geometry only | 0.5273 |
| ESM-2 + confidence | 0.5822 |
| ESM-2 + geometry | 0.5369 |
| Confidence + geometry | 0.5684 |
| Full TAPAS | **0.5847** |

**Supplementary Table S7. Comparison of classifiers on external benchmarks (ePytope-TCR, IMMREP25).** Performance of TabPFN is compared with different classifiers: random forest, XGBoost, a multilayer perceptron (MLP), and logistic regression. Each value is the macro-AUC@0.1 of the corresponding 10-model ensemble.

a. ePytope-TCR

| Model | macro-AUC@0.1 |
| --- | --- |
| TabPFN | **0.5618** |
| Random forest | 0.5578 |
| XGBoost | 0.5608 |
| MLP | 0.5537 |
| Logistic regression | 0.5465 |

b. IMMREP25

| Model | macro-AUC@0.1 |
| --- | --- |
| TabPFN | **0.5847** |
| Random forest | 0.5767 |
| XGBoost | 0.5827 |
| MLP | 0.5541 |
| Logistic regression | 0.5684 |

**Supplementary Table S8. Computational cost of AlphaFold3 structure generation and downstream TAPAS processing for IMMREP25.** AlphaFold3 wall time represents elapsed runtime under the stated degree of parallelization. Five structural samples were generated per pair, and only the top-ranked structure was used for downstream processing. Hardware: 1 × NVIDIA RTX 4500 Ada Generation GPU, Intel(R) Xeon(R) w3-2535 CPU (10 cores / 20 threads).

| Dataset | AF3 MSA and structure prediction  (per TCR-pMHC pair) | Feature extraction  (per TCR-pMHC pair) | TAPAS fitting and inference  (All 10 ensemble models) |
| --- | --- | --- | --- |
| IMMREP25 | 780 sec | 12 sec | 151 sec |
